# PoLoCo: A reproducible pooled low-coverage workflow for draft genome assembly and allele frequency analysis from ethanol-preserved small non-model invertebrates

**DOI:** 10.64898/2026.09.19.752856

**Authors:** Mohammad Jamil Shuvo, Gernot Segelbacher, Julia C. Geue

## Abstract

**Background:** Many ecologically important small invertebrates are preserved in ethanol, which compromises DNA integrity and severely limits their potential to be utilized in further genomic research. The lack of soil arthropod representation ultimately results in limited availability of reference genomes and population genomic inference from pooled sequencing (Pool-seq).

**Results:** We introduce PoLoCo (Pooled Low-Coverage), a reproducible workflow integrating pooled specimen draft genome assembly from ethanol-preserved specimens with population-level allele-frequency analysis. We assembled a 411 Mb draft genome of *Entomobrya nivalis* from pooled short reads, achieving 97.7% BUSCO completeness despite notable fragmentation. Comparison with the published reference genome showed partial genomic correspondence while also revealing reference-dependent differences in the retained SNP datasets. The application of PoLoCo to 82 retained pooled populations produced a final distance-thinned allele-frequency dataset suitable for population-scale genomic inference. Case-study execution documented the computational requirements of the major workflow components on standard HPC infrastructure, with the workflow, documentation, and software environments openly available in Zenodo.

**Conclusions:** PoLoCo demonstrates that draft genome assemblies and reproducible Pool-seq allele-frequency analyses can be performed with long-preserved sample material from the field. PoLoCo integrates reference construction, sync-based allele-frequency analysis, and benchmarking into a free and open workflow. It establishes a reproducibility benchmark for biodiversity genomics and yields a transferable workflow for ecological and evolutionary initiatives where high-quality DNA or long-read sequencing is not feasible.

## 1. Introduction

Recent advances in biodiversity and ecological genomics are making genome-wide data more readily available to characterize population structure, adaptive variation, and evolutionary responses across diverse taxa. Depending on species and context, obtaining such information from non-model and small-bodied taxa often requires scalable, cost-effective sequencing approaches capable of producing genome-wide allele frequency patterns across many populations. This is largely because small-bodied taxa typically yield limited amounts of DNA per individual, while robust population genomic inference depends on sampling many individuals across many populations, making individual-based whole-genome sequencing logistically and financially suboptimal [1–3]. Consequently, pooled sequencing (Pool-seq) has become a widely used and cost-effective approach for population genomic studies, particularly when projects involve many populations or organisms for which individual sequencing is impractical, and is now routinely applied to investigate population structure, local adaptation, demographic patterns, and genotype-environment associations in non-model taxa [1–4]. In practice, combining DNA from several individuals into a pooled library gives a straightforward way to estimate allele frequencies across populations without sequencing each specimen on its own [1,4]. However, even with this efficiency and utility, Pool-seq critically depends upon the availability of an appropriate reference genome. Pool-seq requires good read alignment, which is strongly influenced by reference genome quality and can hinder or even influence the discovery and interpretation of genomic variation [5,6].

This dependency on a high-quality reference genome poses a real challenge for many small-bodied invertebrates. Though ecologically diverse and abundant in nature, they are still underrepresented in genomic databases and other large-scale sequencing initiatives, including community-driven initiatives such as the i5K Consortium [7] and MetaInvert [8], as well as broad biodiversity efforts such as the Earth BioGenome Project [9] and the European Reference Genome Atlas [10], which focus on expanding genomic resources for invertebrate taxa through coordinated community-based efforts. For many of these taxa, a closely related reference genome simply does not exist, and producing one is not straightforward. Small arthropods usually provide only small amounts of DNA, and specimens collected during ecological surveys are often stored in ethanol for later processing. While ethanol is convenient for sampling and storing in various environments, it also can fragment DNA and introduce chemical modifications [11,12]. Recent advances in museomics show however, that some ethanol-preserved collection material retain sufficiently long DNA fragments which can support long-read genome assembly [13,50]. In the case of field-preserved specimens, with more fragmented or low-input DNA, however, short-read sequencing remains still a practical alternative, although assemblies generated from such material may have limited contiguity [13,14]. These difficulties become even more noticeable when the material comes from pooled specimens, as they tend to show higher heterozygosity and often lead to assemblies with redundant or less contiguous regions, which complicates read mapping and downstream variant calling [15].

These limitations affect the entire Pool-seq workflow, because using an incomplete or distant reference introduces systematic alignment bias. Such bias may inflate apparent rare variants, distort allele frequency distributions, cause mapping failures in divergent regions, and generate local clusters of spurious SNPs [16,17]. Although established Pool-seq tools such as PoPoolation2 [4], SNAPE [18], and PoolParty [2] provide useful frameworks for pooled allele-frequency analysis, they assume that a suitable reference genome is already available. As a result, key components of the Pool-seq workflow, including reference genome construction, assembly validation, and the handling of low-input or degraded DNA, are typically addressed outside these tools. Consequently, existing pipelines do not offer an integrated solution for studies in which long-read sequencing is not feasible, and population-level analyses must rely on degraded or low-quantity material. This gap limits the application of Pool-seq to many ecologically important systems where high-quality DNA cannot be obtained.

To address this need, we developed PoLoCo (Pooled Low-Coverage), a reproducible workflow designed to integrate draft genome assembly, reference validation, and population-level allele-frequency estimation to enable Pool-seq analyses from ethanol-preserved, pooled invertebrate specimens and other non-model taxa where high-quality DNA or long-read data are not available. Rather than introducing a new genome assembler or allele-frequency estimator, PoLoCo addresses the separation between project-specific reference generation and downstream Pool-seq analysis, which are typically handled as independent stages. PoLoCo connects draft reference construction and validation directly with configurable read mapping, sync-based SNP filtering and allele-frequency estimation, quality control, and reproducible execution in a single workflow. In the first step of the PoLoCo workflow, a draft genome can be assembled by pooling short-read sequences, which provides a reference assembly suitable for downstream analyses. The draft reference is then validated using available genomes. The validated draft genome is subsequently used as the reference for the Pool-seq analyses, including quality control, read mapping, sync-based SNP filtering, and estimation of allele frequencies. PoLoCo is distributed with a single installation script for its modular Conda environments, centralized user-configurable settings, and sequential or dependency-aware SLURM launchers for reproducible execution.

Because springtails (Collembola) are small-bodied invertebrates, routinely preserved in ethanol, and rarely have high-quality reference genomes, they offer a realistic system for evaluating workflows designed for degraded, low-input material. As a case study, we applied the PoLoCo workflow to the springtail *Entomobrya nivalis*, a widespread species in temperate forest soils. Collembola play an important role in soil ecosystems by contributing to nutrient cycling, decomposition, and the regulation of microbial activity [19–21]. Despite their ecological importance, genomic resources for Collembola remain scarce compared with those available for many insects and vertebrate taxa [22,23]. Our work had four main aims. First, we assembled a draft genome of *E. nivalis* from pooled ethanol-preserved specimens using short-read data. Second, we compared this draft genome assembly with the published *E. nivalis* genome to evaluate its accuracy. Third, we tested how well the draft genome supported SNP identification and allele-frequency estimation across a final dataset of 82 pooled populations. Finally, we assessed the computational resources needed to run the workflow so that it can be reproduced and scaled on typical high-performance computing systems.

## 2. Materials and Methods

### 2.1. Sample collection and DNA extraction

*Entomobrya nivalis* specimens were collected as part of the ConFoBi (Conservation of forest biodiversity in multiple-use landscapes of central Europe) Research Training Group sampling effort in 2017. The ConFoBi project has 135 one-hectare plots established since 2016 in the southern Black Forest, Germany. The area spans >5,000 km² of mixed forests dominated by *Picea abies*, *Fagus sylvatica*, *Abies alba*, and other common species, including oak, maple, and Douglas fir. Plots range from 443 to 1,334 m elevation with slopes up to 34° [24]. The Collembola specimens were collected from flight interception traps (FIT), which were primarily set up to monitor flying arthropods. In total, 9,457 individual Collembola specimens were recovered across all 135 plots and sorted to at least family-level, based on their morphology. All specimens were placed in 99% molecular-grade ethanol immediately after collection and stored at −20 °C in the specimen archive of the Chair of Wildlife Ecology and Management at the University of Freiburg. Due to the small body size of Collembola and ethanol storage causing DNA fragmentation, DNA extraction from single specimens did not yield enough high-molecular-weight DNA for whole-genome sequencing. DNA was pooled prior to extraction to yield sufficient DNA for pooled sequencing. DNA extraction was performed using the Qiagen DNeasy Blood & Tissue Kit (Qiagen, Hilden, Germany). Whole bodies were used for the extraction, and the lysis step was extended overnight at 56 °C with proteinase K to maximize DNA yield from chitinous tissue. DNA was purified and concentration determined with the Qubit dsDNA HS Assay Kit on a Qubit 3.0 or 4 Fluorometer (Thermo Fisher Scientific). For draft genome assembly, a single pooled sample consisting of eight adult *E. nivalis* individuals was selected to maximize sequencing depth. To apply the PoLoCo workflow at the population scale, we initially prepared 93 additional pooled samples, each consisting of 6-10 individuals collected from a single ConFoBi plot. After quality control and downstream filtering, 82 pooled libraries were retained and used to evaluate the performance of the workflow for population-level analyses using low-coverage ethanol-preserved material (see Sections 3.1.2 and 3.3).

### 2.2. Library preparation and sequencing

Whole-genome libraries were constructed using the Watchmaker Genomic DNA Library Prep Kit (Watchmaker Genomics, USA) due to its ability to utilize degraded and low-input DNA, such as ethanol-preserved material. Libraries were constructed from pooled DNA while retaining shorter DNA fragments to accommodate the fragmented nature of the ethanol-preserved material. For the draft genome assembly, we selected one pooled library representing a typical ConFoBi plot. This pool was chosen based on its high DNA yield and overall DNA integrity relative to other pools, ensuring sufficient input for library construction. The selected library was sequenced to approximately 25× coverage. The remaining pooled population libraries were sequenced at Floragenex (currently part of Rapid Genomics, USA) using the Illumina NovaSeq 6000 platform, with SP 300-cycle flow cells to generate 150 bp paired-end reads. The library kit and sequencing strategy were chosen to ensure sufficient data yield from degraded ethanol-preserved material and to allow the workflow to scale to a large population dataset. All raw sequence data were stored in compressed FASTQ files for preprocessing and quality control.

### 2.3. Read preprocessing and quality control

The preprocessing was done on all raw sequence data prior to any downstream analysis, which included removing adapters, filtering low-quality bases, and generating quality reports. Preprocessing was done using fastp v0.23.2 [25], which is a multi-purpose FASTQ preprocessor and suitable for large Illumina datasets. The application was used with default parameters, and specified filters to retain sequences that had Phred scores of Q20 or greater, and removed reads that had less than 50 bp of remaining length, after trimming. Fastp auto-detected and trimmed the adapter sequences. Each pooled population library (the assembly dataset was handled separately) was processed individually and stored in gzip-compressed FASTQ files for downstream processing. Read quality was assessed at two time points, first on the raw FASTQ files and again after preprocessing to verify that filtering had been applied as intended. Quality reports were generated before and after preprocessing using FastQC v0.11.9 [26] and collated using MultiQC v1.14 [27]. These included standard summaries of read quality, adapter content, and sequence length distributions. All preprocessing steps were performed in an automated fashion using the PoLoCo workflow, a reproducible pipeline integrating read preprocessing, draft genome assembly, and validation, and population-level Pool-seq analyses, implemented through dedicated conda environments and SLURM batch scripts to ensure reproducibility across all datasets. The full workflow and scripts are available in Zenodo [28].

### 2.4. Genome assembly and validation

For de novo assembly of the *Entomobrya nivalis* draft genome, we used a high-coverage pooled dataset. We assembled the data with MEGAHIT v1.2.9 [29], a de Bruijn graph-based assembler intended for large and complex short-read datasets such as those obtained from degraded DNA and pooled individuals. The assembler was run using the --meta-sensitive preset to optimize graph construction, and k-mer sizes were automatically selected in the range of 21 to 119. After assembly, contigs less than 1,000 bp were removed in order to remove the noise from spurious assemblies and very short fragments that may be uninformative for downstream analyses. The filtered assembly was used for all subsequent analyses. Assembly statistics including total assembly size, number of contigs, N50, and L50 were calculated using QUAST v5.2.0 [30]. Assembly completeness was quantified using BUSCO v5.5.0 [31] in genome mode using the arthropodaodb10 lineage dataset (1,013 single-copy orthologs). Comparative validation was carried out by aligning the draft genome assembled in this study against the previously published *E. nivalis* reference genome (NCBI access GCA_034695485.1). Whole genome alignments were done using MUMmer4 v4.0.0 [32], and dot plots were generated using the mummerplot utility included in the MUMmer package. Dot plots visualize pairwise genome alignments by plotting matching regions between the draft and reference genomes, allowing assessment of large-scale structural similarity, collinearity, and potential misassemblies. Assembly correspondence was further quantified using the MUMmer4 dnadiff [32] workflow, which reports the fraction of each assembly represented in the alignment together with nucleotide identity across aligned regions. We used the dnadiff one-to-one alignment summary as the primary quantitative sequence-comparison metric. FastANI v1.33 [33] was retained as an auxiliary diagnostic and was not used to infer biological divergence between the conspecific assemblies. QUAST v5.2.0 was run in comparison mode, producing metrics such as genome fraction and misassembly counts that rely on alignment.

### 2.5. Mapping of pool-seq data

We aligned the retained pool-seq libraries to the *E. nivalis* draft genome produced in this study using BWA-MEM v0.7.17 [34], which is suited for Illumina short reads and tuned for gapped alignments. For reference-performance benchmarking, the same retained pooled libraries were also aligned to the published *E. nivalis* reference genome (GCA_034695485.1) and processed through the same downstream Pool-seq workflow. We aligned each pooled dataset independently, creating a sequence alignment map (SAM) file as the initial output. Alignments were processed using samtools v1.17 [35] to generate sorted and indexed Binary Alignment/Map (BAM) files. Duplicated reads were removed using Picard MarkDuplicates v2.26.10 [36]. Subsequently, only properly paired alignments with a mapping quality of at least 30 were retained, and mpileup generation was performed with a minimum base quality threshold of 20. Both parameters are specified in the central PoLoCo configuration and may be adjusted according to the requirements of individual datasets. Coverage per base was estimated using samtools depth, and summary statistics (mean depth, mapping rate) were generated with Python scripts in the PoLoCo repository. All steps were run as modular Slurm batch scripts in conda environments to ensure versions are consistent and reproducible.

### 2.6. SNP discovery and allele frequency estimation

Single nucleotide polymorphisms and pool-level allele frequencies were derived from a synchronized pooled read-count file generated from filtered BAM alignments using samtools mpileup and the PoPoolation2 utility mpileup2sync.pl [4,35]. Downstream SNP filtering, allele-frequency extraction, and matrix construction were then performed from this sync file using a reproducible shell-based Bash pipeline implemented in PoLoCo. This downstream procedure followed the general count-based logic of pooled allele-frequency analysis from synchronized read-count data as used in PoPoolation2 and related Pool-seq studies [4,16,17], while the specific filtering and matrix-building steps were implemented as part of the present workflow. To restrict the analysis to true polymorphic sites, nucleotide counts for A, T, C, and G were first summed across all pools at each genomic position, and only positions with at least two alleles showing non-zero total counts were retained for further processing. Pool-level allele frequencies were then estimated directly from nucleotide read counts rather than from genotype calls. For each pool at each retained site, coverage was defined as the sum of A, T, C, and G counts. A pool was considered informative at a site only when coverage was at least 4 reads. Sites were retained only if they were represented in at least 55 of the 82 pooled populations (at least 67%), corresponding to approximately two thirds of all retained pools. This threshold was chosen to balance locus retention against missingness, so that downstream analyses were based on SNPs represented across a substantial proportion of populations while still allowing some low-coverage pools to be missing at individual sites. These values represent the settings used for the *E. nivalis* case study. In the PoLoCo workflow, the minimum coverage, minimum population representation, minor-allele frequency, and thinning distance are user-configurable, and population representation can be specified either as an absolute number of pools or as a proportion of the dataset (e.g., 55 pools or a proportional threshold of 0.67 for the present 82-pool case study). For every retained site, total nucleotide counts were summed across all informative pools, and the globally most abundant allele was assigned as the major allele whereas the second most abundant allele was assigned as the minor allele. The global minor allele frequency was calculated as the total read count of the minor allele divided by the summed read count across all informative pools. For each population, the minor-allele frequency used in downstream analyses was then calculated as the read count of the globally defined minor allele divided by the total site coverage in that pool. Pools failing the minimum coverage threshold at a given site were treated as missing values for that site. To obtain the final analytical dataset, SNPs were filtered to retain only loci with a global minor allele frequency of at least 0.05. Marker redundancy among nearby sites was then reduced by simple distance-based thinning along each scaffold, using 200 bp as a pragmatic case-study spacing criterion rather than as an estimate of linkage-disequilibrium decay. SNP identifiers were decomposed into scaffold and genomic position, loci were ordered by coordinate within scaffold, and the first SNP encountered on each scaffold was retained. Each subsequent SNP was retained only if it occurred more than 200 bp away from the previously retained SNP on the same scaffold. Final outputs consisted of a SNP annotation table containing scaffold, position, major allele, minor allele, and the number of informative pools, together with a SNP-by-population minor-allele-frequency matrix used for downstream analyses.

### 2.7. Workflow benchmarking, reproducibility, and data availability

An overview of the PoLoCo workflow, including separate assembly and population inputs, the ordered processing steps, QC checkpoints, and the optional population-structure and spatial-connectivity analysis, is shown in Figure 1. Workflow performance was assessed on bwUniCluster 3.0 HPC systems. The core workflow (Steps 0-7; Figure 1) can be executed sequentially using the master Bash runner script (scripts/run_poloco_pipeline.sh) or submitted as dependency-aware SLURM jobs, in which downstream jobs are released only after successful completion of their required preceding steps. Before execution, the input-check and setup step (Step 0; Figure 1) verifies configured inputs, paired-read structure, reference availability, and core analysis parameters before computationally intensive steps are initiated. The optional final step (Step 8: population-structure and spatial-connectivity analysis; Figure 1) uses the workflow-generated allele-frequency matrix to perform principal component analysis (PCA), calculate pairwise genetic and geographic distances, and evaluate isolation by distance using a Mantel test with 9,999 permutations. Projected plot coordinates are supplied separately and matched to the population order recorded by the workflow. This module is intended as a concise demonstration of downstream use of PoLoCo output rather than as a comprehensive population-genomic analysis. No single step used more than 32 CPU cores or 128 GB RAM. Runtime and memory limits for each step are listed in Results (Section 3.3) and provided in Table 1. Raw Illumina reads from the assembly dataset and the pooled population libraries retained for downstream analyses are available in the European Nucleotide Archive (ENA) under BioProject accession PRJEB111482 [37]. The draft genome assembly, BUSCO outputs, QUAST reports, and all supplementary files are stored on Figshare [38], along with a mirrored copy of the assembly archived in ENA. The full PoLoCo workflow, modules, conda environments, SLURM scripts, and QC utilities are available in Zenodo [28].

**Figure 1.**
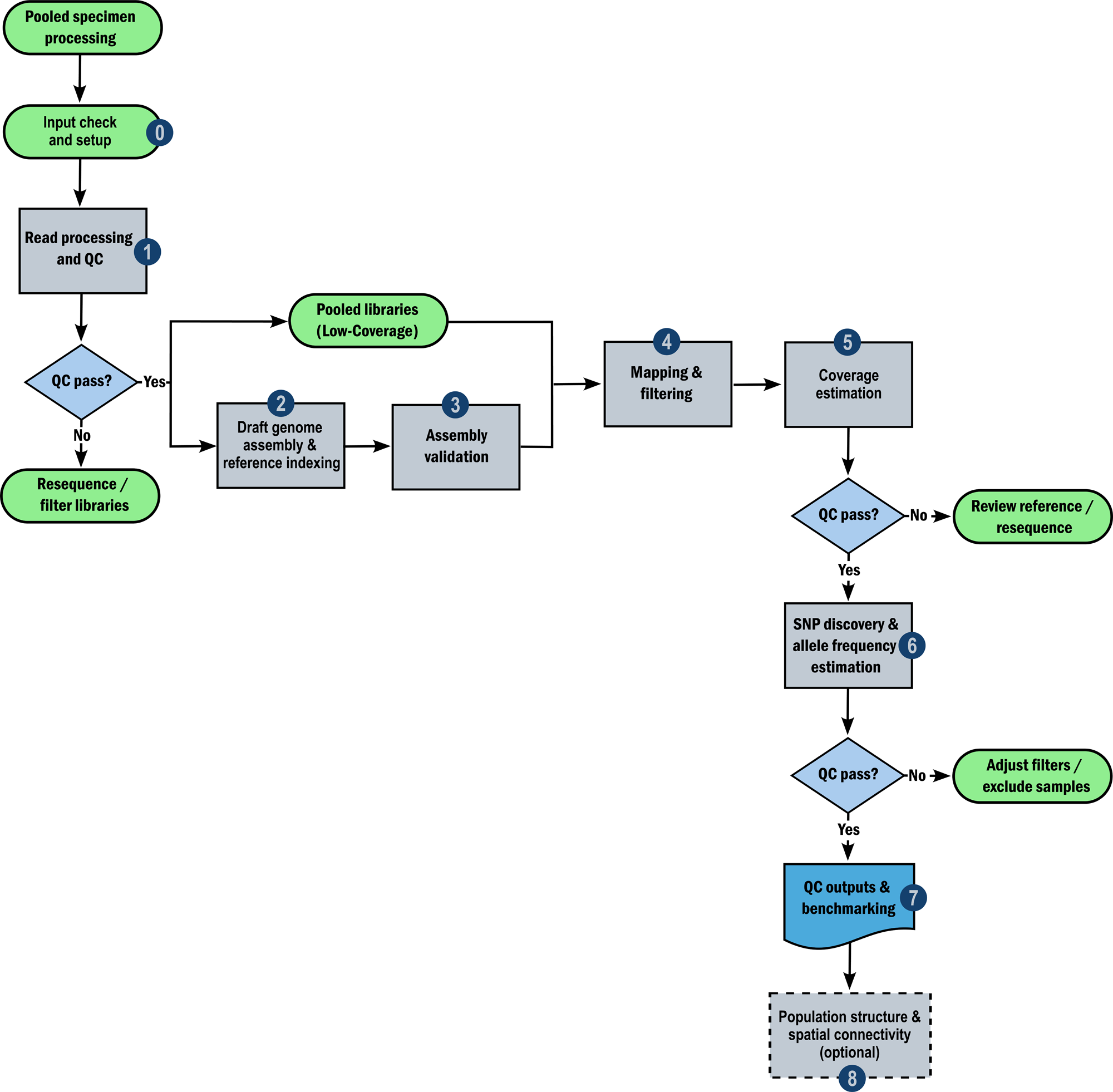
PoLoCo workflow diagram. Rounded rectangles denote processing steps, diamonds denote QC checkpoints, ovals denote potential outcomes/checkpoints, and dashed boxes denote optional modules. Solid arrows indicate the primary data flow, whereas side arrows indicate alternative routes when QC checkpoints fail. Numbered labels 0-8 correspond to the ordered workflow scripts in the GitHub repository. Following input check and setup (Step 0) and read processing and QC (Step 1), the workflow separates into a draft-genome branch and a pooled low-coverage population-library branch. The draft genome is assembled and indexed as the mapping reference (Step 2) and then validated (Step 3) before the pooled libraries are mapped and filtered (Step 4). Subsequent steps comprise coverage estimation (Step 5), SNP discovery and allele-frequency estimation (Step 6), QC outputs and benchmarking (Step 7), and the optional population-structure and spatial-connectivity analysis (Step 8). **Alt text:** Workflow diagram showing the PoLoCo pipeline from input validation and read preprocessing through draft genome assembly and validation, pooled-library mapping and filtering, coverage estimation, SNP discovery and allele-frequency estimation, QC and benchmarking, and optional population-structure and spatial-connectivity analysis. Numbered labels 0-8 correspond to the ordered GitHub workflow scripts.

**Table 1.** Runtime and memory requirements for PoLoCo workflow steps.

| Step | Tools | Typical Runtime | CPUs | RAM | Outputs |
| --- | --- | --- | --- | --- | --- |
| Preprocessing<br>& QC | fastp, FastQC, MultiQC | 1-2 h / pool | 4 | 32 GB | Trimmed reads, QC reports |
| Assembly | MEGAHIT, BUSCO | ~48 h | 16 | 128 GB | Draft genome assembly |
| Validation | QUAST, MUMmer4/dna diff, FastANI | 2-3 h | 8 | 64 GB | Assembly metrics, aligned-region identity, reference-comparison outputs |
| Mapping & Filtering | BWA, samtools, Picard | 2-3 h / pool | 8 | 64 GB | Deduplicated & filtered BAMs |
| Coverage Estimation | samtools depth | <1 h | 4 | 16 GB | Coverage table per pool |
| Pool-seq pipeline | samtools, PoPoolation2, Bash workflow | 6-8 h (20 pools); ~24 h (82 pools) | 8-16 | 64 GB | mpileup, sync file, filtered SNP table, allele-frequency matrix |
| Population structure & connectivity | R, PCA, vegan | ~1 min | 2 | 16 GB | PCA, spatial map, pairwise distances, Mantel test |

Benchmarks were obtained on bwUniCluster HPC nodes as described in Methods (Section 2.7). Estimates are based on pooled *Entomobrya nivalis* Illumina short-read datasets (20-82 pools; approximately 7-15 million reads per pool) run on standard HPC nodes. Reported values summarize the typical runtime, computational resource allocations, and principal outputs for each step in the case-study execution.

## 3. Results

### 3.1 Read quality and filtering of pooled samples

#### 3.1.1 Draft genome assembly, completeness, and validation

The draft genome assembly of *Entomobrya nivalis* utilized a library of pooled DNA from eight individuals and yielded high read depth for assembly, corresponding to approximately 25× genome coverage. Fastp quality control indicated that the usable data retention was high, with 94.2% of the raw reads retained after filtering. The per-base quality scores were consistently high throughout the dataset, with 96% of bases exceeding Q20 and 95% of bases exceeding Q30. Although DNA was degraded, short-read sequencing produced data of sufficient quality for assembly. The GC content of the assembly was tightly clustered at 38.0%, which is not unexpected for Collembola genomes and is consistent with the GC content reported for the published *Entomobrya nivalis* reference genome [39]. No adapter contamination was detected (<0.1%), and duplication rates were <2%. Figure 2a summarizes the assembly dataset QC, showing GC content, Q30 values, and read retention for the pooled library used for assembly. These metrics confirm that the dataset had an expected GC content (38%), high per-base quality, and strong read retention (>94%). Together, these results indicate that the sequencing data were suitable for producing a reliable de novo genome assembly. After discarding contigs < 1 kb, the assembly contained 142,936 contigs and a total length of 411.39 Mb (Table 2). The longest contig was 463,496 bp, while the fragmented nature of the assembly was reflected by an N50 of 3,617 bp and an L50 of 30,950 contigs. Cumulatively, contig size distribution indicates that while the majority of assembled fragments were shorter, there was a sizable number of longer scaffolds recovered. The assembly had 3,622 contigs with ≥10 kb, 205 contigs with ≥25 kb, and 36 contigs with ≥50 kb. The reference values were predictably expected from an assembly from short-read Illumina sequencing of DNA, which was in ethanol for a prolonged amount of time.

**Figure 2.**
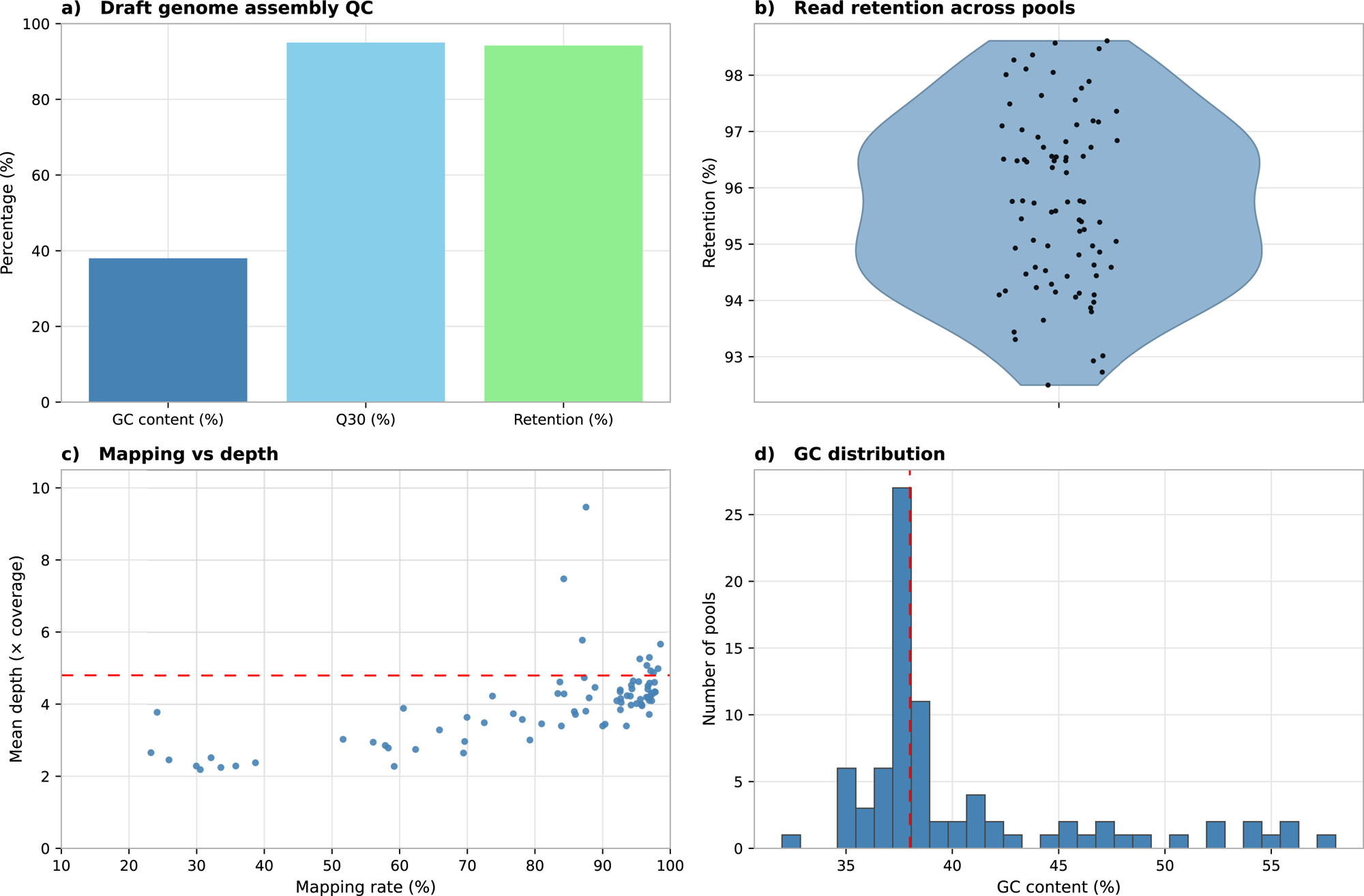
Quality and filtering metrics for assembly and pooled libraries. (a) Bar plot showing quality statistics for the assembly dataset derived from pooled individuals used to generate the draft genome, including GC content (blue), Q30 (light blue), and read retention (green). (b-d) Quality metrics for the pooled population libraries retained for the final Pool-seq analysis. (b) Violin plot of read retention (%) across pooled samples; black dots represent individual samples. (c) Scatter plot of mapping rate (%) versus mean sequencing depth (× coverage) across pooled samples; each blue point represents one pooled sample. The dashed red line indicates the fitted linear regression. The y-axis is capped at 0-10× coverage to highlight the main cluster; a few samples exceeded this range. (d) Histogram of GC content (%) across pooled samples; the x-axis shows GC percentage and the y-axis shows number of pooled samples. The dashed red vertical line indicates the expected mean GC content (∼38%). **Alt text:** Four-panel figure summarizing sequencing quality metrics for the assembly dataset and retained pooled population libraries, including GC content, Q30, read retention, mapping rate versus depth, and GC-content distribution.

**Table 2.** Assembly and pairwise comparison metrics for the draft and published reference genomes.

| Metric | Draft genome (this study) | NCBI reference<br>(GCA_034695485.1) |
| --- | --- | --- |
| Total assembly size (Mb) | 411.4 | 300.9 |
| Contigs ($\geq 1$ kb) | 142,936 | 41,156 |
| N50 (bp) | 3,617 | 15,827 |
| L50 (contigs) | 30,950 | 3,000 |
| Largest contig (bp) | 463,496 | 128,347 |
| GC content (%) | 37.4 | 34.5 |
| Genome fraction (%) | 97.9 | 100 |
| Misassemblies | 62 | - |
| Assembly aligned (%) | 29.58 | 38.65 |

Misassemblies are QUAST reference-based counts for the draft assembly evaluated against GCA\_034695485.1. The published NCBI assembly was supplied as the reference and was not evaluated against itself. Therefore, no independent misassembly count is reported for the NCBI-reference column.

Overall GC content of the assembly equals 37.4%, within the reported ranges of Collembola genomes and the GC composition of the raw reads. This is further verification that the assembly did not have systematic compositional bias. Completeness of the genome was evaluated with BUSCO v5.5.0 using the arthropodaodb10 lineage dataset of 1,013 conserved orthologs. The analysis reported that there were 990 complete BUSCOs (97.7%), 521 single-copy BUSCOs, and 469 duplicated BUSCOs. There were 12 BUSCOs (1.2%) that were fragmented, and 11 missing BUSCOs (1.1%) (Table 3). These values indicate a high level of BUSCO completeness for the assembly. Figure 3 illustrates the draft genome assembly statistics and alignment results between the PoLoCo draft genome assembly and the published *E. nivalis* reference genome (GCA_034695485.1). To validate the genome assembly, we performed comparative alignments with the published *E. nivalis* reference genome. Whole-genome comparison with the published *E. nivalis* reference genome showed that the draft assembly recovered 97.9% of the reference genome fraction (Table 2). QUAST comparison also identified 62 misassemblies in the draft assembly, which is expected for a short-read assembly generated from degraded pooled material and reflects its fragmented structure relative to the more contiguous published reference. MUMmer4 dnadiff identified one-to-one aligned regions with an average nucleotide identity of 87.96%. Aligned bases represented 38.65% of the published reference and 29.58% of the draft assembly. The Nx cumulative length curve (Figure 3a) is presented to show the distribution of fragment size with the N50 of 3,617 bp marked. Visualization with MUMmer4 showed a diagonal trend among aligned regions in the dot plot, although substantial portions of both assemblies remained unaligned (Figure 3b). The comparison therefore indicates partial genomic correspondence rather than near-complete collinearity between the assemblies. Collectively, these provide a draft genome assembly that was used later for SNP discovery and estimation of allele frequency.

**Figure 3.**
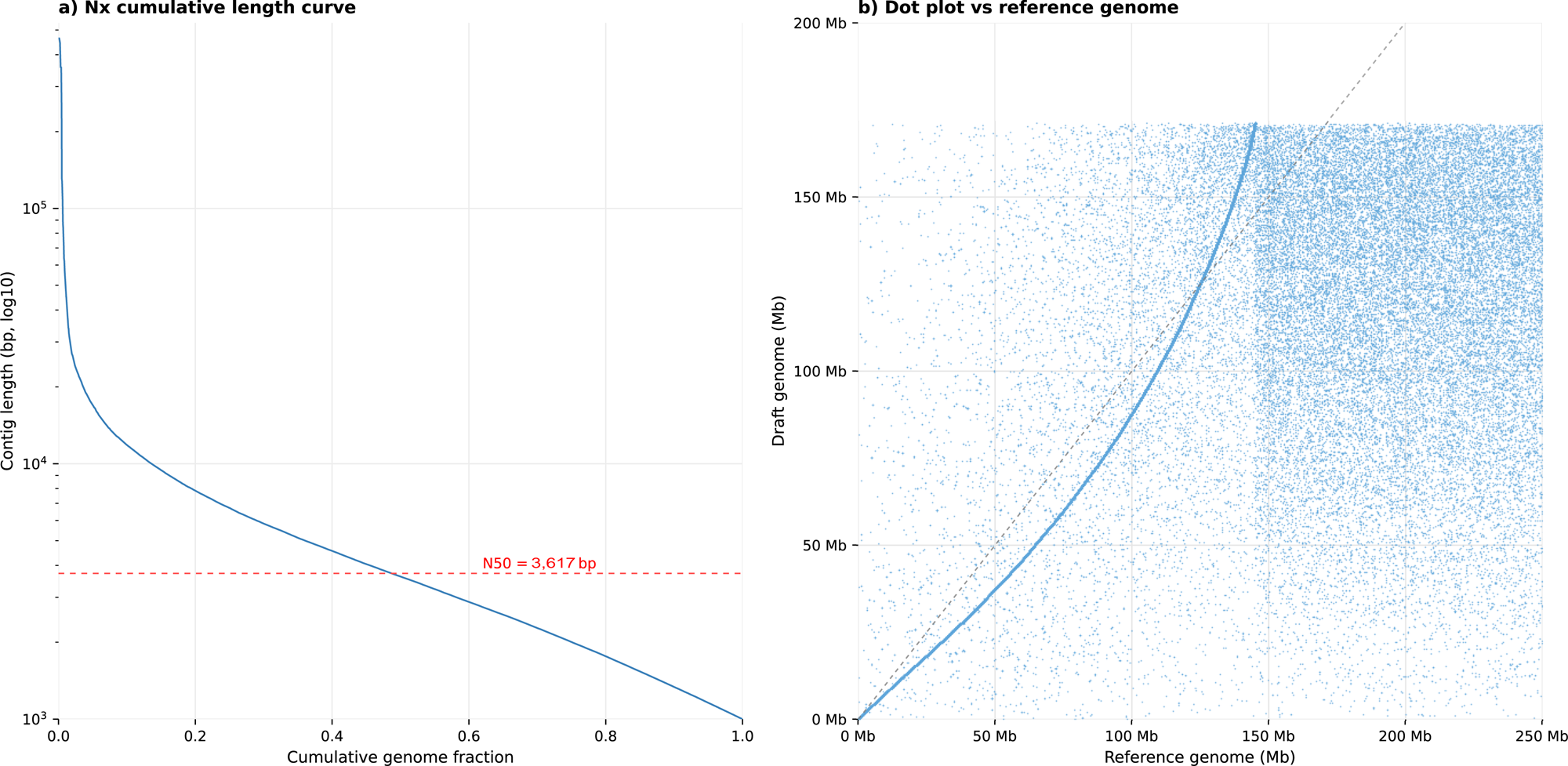
Draft genome assembly statistics and reference alignment. (a) Nx cumulative length curve for contigs ≥1 kb; the x-axis shows cumulative genome fraction and the y-axis shows contig length in log10 scale (bp). The N50 value is shown as a red dashed horizontal line at 3,617 bp. (b) Dot plot comparing the draft genome assembly with the published reference genome (GCA_034695485.1); each dot indicates an aligned region, and the axes show genome coordinates in megabases (Mb). The diagonal dashed line represents perfect collinearity. **Alt text:** Two-panel figure showing draft genome assembly evaluation, including an Nx cumulative contig-length curve with N50 marked and a dot plot showing diagonal trend among aligned regions, together with substantial off-diagonal and unaligned sequence.

**Table 3.** BUSCO assessment of the draft *Entomobrya nivalis* genome.

| Category | Count | Percentage |
| --- | --- | --- |
| Complete BUSCOs (C) | 990 | 97.7% |
| Single-copy (S) | 521 | 51.4% |
| Duplicated (D) | 469 | 46.3% |
| Fragmented (F) | 12 | 1.2% |
| Missing (M) | 11 | 1.1% |
| **Total** | **1013** | **100%** |

#### 3.1.2 Pool-seq dataset

Quality control was performed on all pooled libraries generated for population-scale analyses, of which 82 were retained in the final dataset. Each library contained between 7.3 and 14.7 million paired-end reads. After quality filtering, retention rates were between 92.5 to 97.1%, with a mean retention above 95% (Figure 2b). Quality metrics were consistently high across the retained libraries: Q20 score was >96% for each pool, and Q30 scores ranged from 90.4 to 94.3%. Mapping quality metrics are shown in Figure 2c, which illustrates the relationship between mapping rate and mean sequencing depth. Mapping rates spanned approximately 23.2-98.5% across the retained pools, with a main cluster at high mapping rates but a subset of libraries below 50%. Most pools clustered within the 2-6 times depth range with a few outliers. Genome representation was high (90% at ≥1× across all pools), which confirmed that the majority of the draft genome was represented in each dataset. GC-content distributions were more heterogeneous than the base-quality metrics: many pools clustered around 37-39% GC, one pool was lower at approximately 32%, and a substantial right tail extended from approximately 40% to 58% GC (Figure 2d). The combination of elevated GC content and reduced mapping in a subset of pools is consistent with the possible presence of residual exogenous sequence in whole-body extractions, although unmapped reads were not taxonomically classified and their origin therefore cannot be determined directly. Such non-target sequence is plausible in genomic data generated from whole-body invertebrate specimens [8]. The conservative alignment and locus-retention criteria used in PoLoCo, including proper-pair and mapping-quality filtering followed by coverage and population-representation thresholds, reduce the influence of poorly supported alignments on the final allele-frequency dataset. Overall, the retained libraries showed consistently high read-retention and base-quality metrics while exhibiting greater variation in mapping performance and GC composition.

### 3.2 SNP discovery and allele frequency estimation

We evaluated the suitability of the *E. nivalis* draft genome assembled in this study for population-scale analyses by conducting SNP discovery and allele-frequency estimation across the final 82 pooled libraries and comparing its performance against the published *E. nivalis* reference genome (GCA_034695485.1). Variant calling was conducted in parallel on both the draft genome assembly and the published NCBI reference genome, and the resulting SNP retention patterns and final allele-frequency datasets were compared. Figure 4 summarizes the filtering outcomes and characteristics of the retained SNP datasets across references. After the minimum informative-population filter, the draft genome yielded a larger number of candidate polymorphic loci than the NCBI reference (Figure 4a). However, after applying the global minor allele frequency threshold and the final 200 bp distance-based thinning step, the NCBI reference retained a substantially larger final SNP set than the draft genome (20,463 vs 5,391 SNPs; Figure 4a). The two final datasets also differed in locus representation across populations. SNPs retained with the draft genome were generally supported by a greater number of informative populations per locus, whereas the NCBI-based dataset contained a larger proportion of loci close to the minimum threshold of 55 informative populations (Figure 4b). Post-filtering allele-frequency distributions also differed between references, with the draft-based dataset showing a stronger enrichment of low-frequency variants and the NCBI-based dataset showing a flatter distribution across the retained minor allele frequency range (Figure 4c). The Ts/Tv ratio further differed between the final retained SNP sets, with a lower value for the draft genome than for the NCBI reference (1.34 vs 1.65; Figure 4d). These values remain within the range generally reported for arthropod genomes [40–42]. Together, these results show that reference choice strongly influenced SNP retention and final dataset composition. The published NCBI reference retained a much larger final SNP set, whereas the project-specific draft genome yielded a smaller but more conservatively retained dataset with stronger population support per locus. This comparison confirms that the draft genome generated in this study performed reliably as a project-specific reference for downstream Pool-seq analyses while also demonstrating the sensitivity of allele-frequency datasets to reference selection. To place this in context, we compare PoLoCo with other popular Pool-seq pipelines (Table 4). The sync-based SNP discovery and allele-frequency estimation presented here were implemented using PoPoolation2 [4] within the broader PoLoCo workflow. Although established tools such as PoPoolation2 [4], ANGSD [43], PoolParty [2], and assessPool [44] provide powerful frameworks for pooled allele-frequency analysis, they assume access to an available reference genome and do not address the challenge of generating project-specific references from degraded material. In contrast, PoLoCo integrates draft reference assembly, reference validation, and downstream PoPoolation2-based sync analysis within a single reproducible workflow, thereby extending Pool-seq applications to ethanol-preserved material.

**Figure 4.**
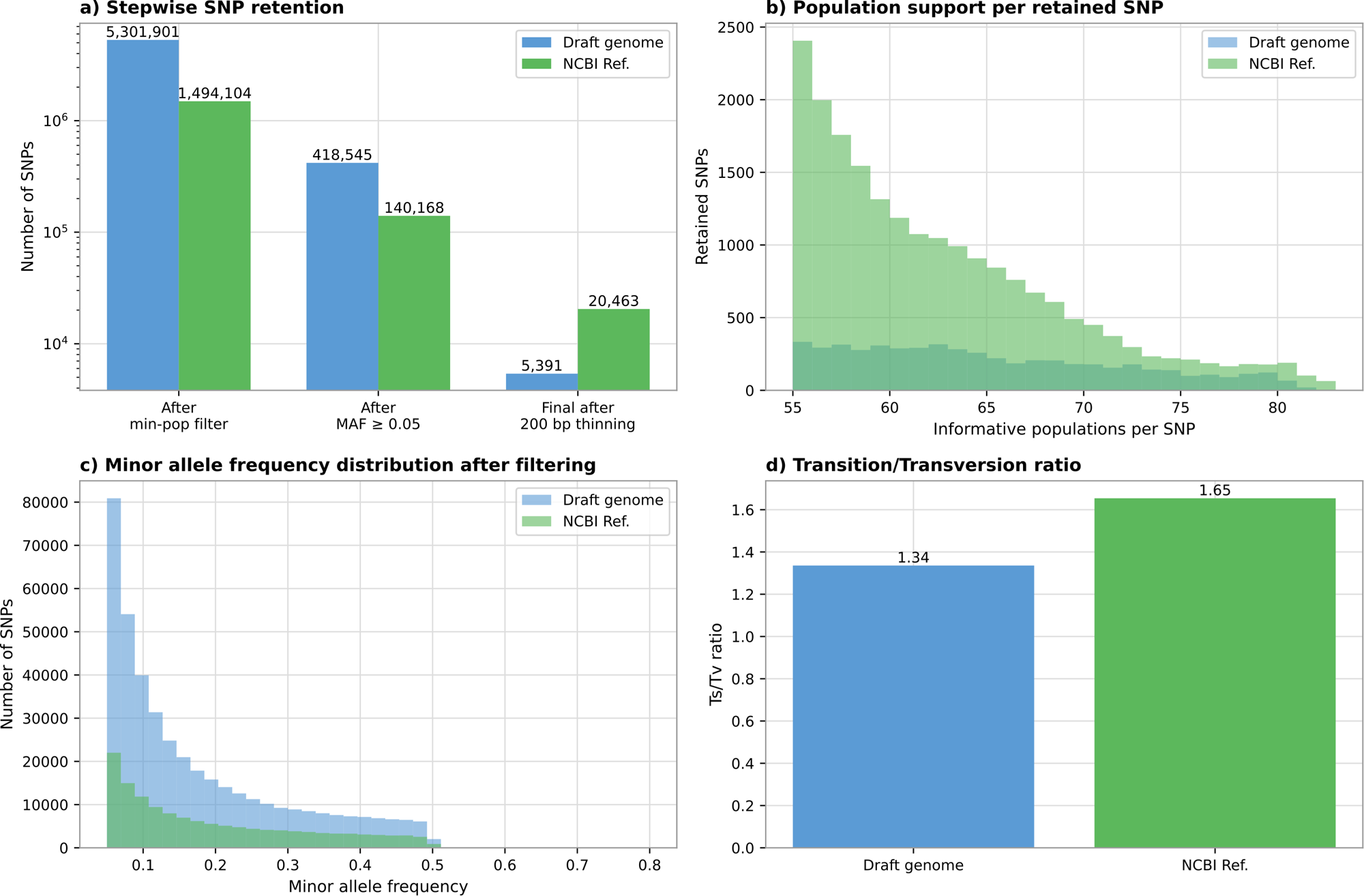
Reference-dependent SNP retention and allele-frequency dataset comparison. (a) Stepwise SNP retention after the minimum informative-population filter, global minor allele frequency filtering, and final 200 bp distance-based thinning. Bars show the number of retained SNPs at each stage for the draft genome (blue) and the NCBI reference (green); the y-axis is shown on a log scale. (b) Distribution of informative populations per retained SNP in the final distance-thinned datasets. The x-axis shows the number of pooled populations with sufficient coverage contributing to each SNP, and the y-axis shows the number of retained SNPs. Colors indicate the draft genome (blue) and the NCBI reference (green). (c) Distribution of minor allele frequencies in the retained SNP datasets after filtering. The x-axis shows global minor allele frequency, and the y-axis shows the number of retained SNPs. Colors indicate the draft genome (blue) and the NCBI reference (green). (d) Transition/transversion (Ts/Tv) ratio of the final retained SNP sets. Bars show the Ts/Tv ratio for the draft genome (blue) and the NCBI reference (green). **Alt text:** Four-panel comparison of SNP filtering outcomes for the draft genome and published reference across 82 pooled populations, including stepwise SNP retention, informative-population support per SNP, minor allele frequency distributions, and Ts/Tv ratios.

**Table 4.**
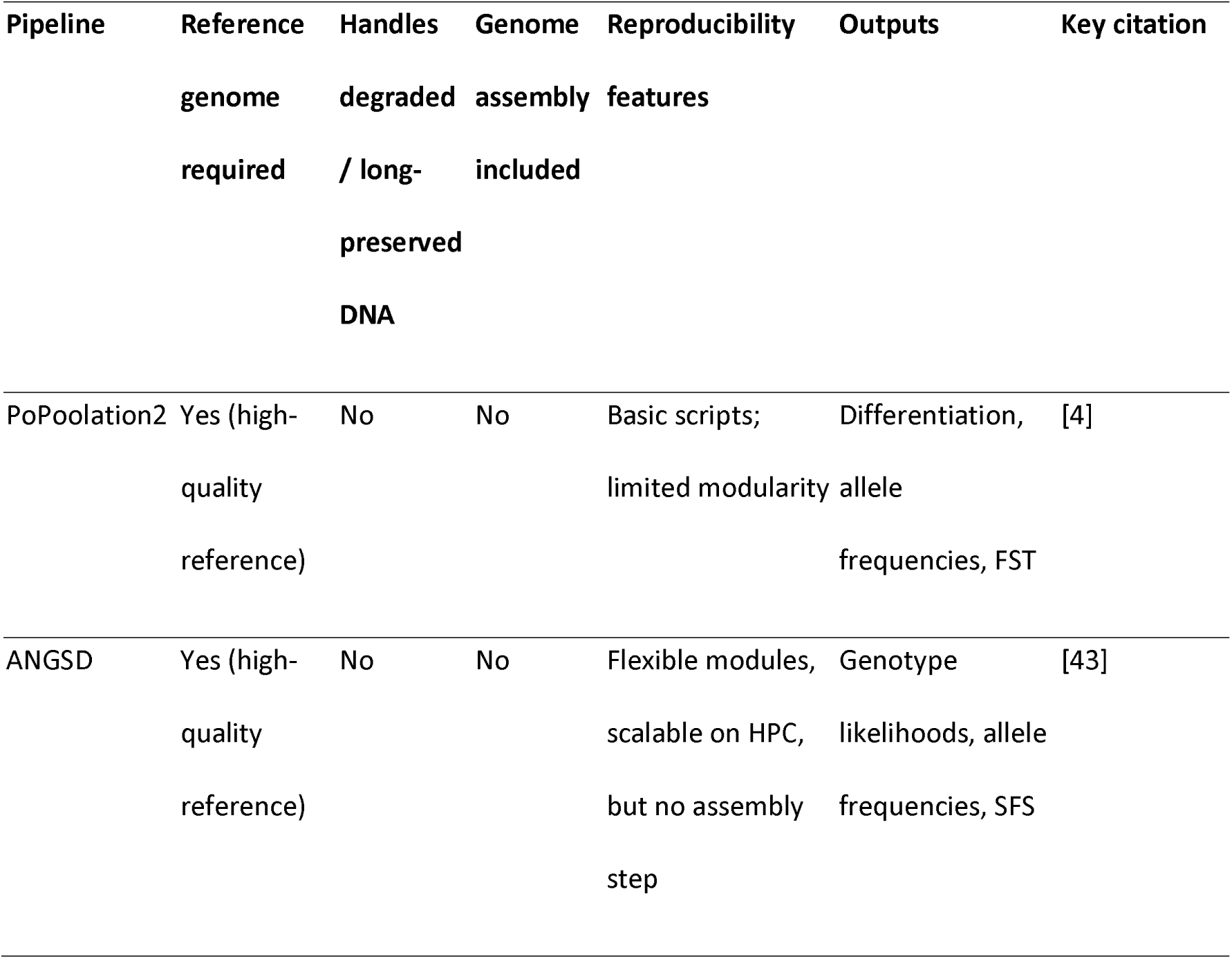
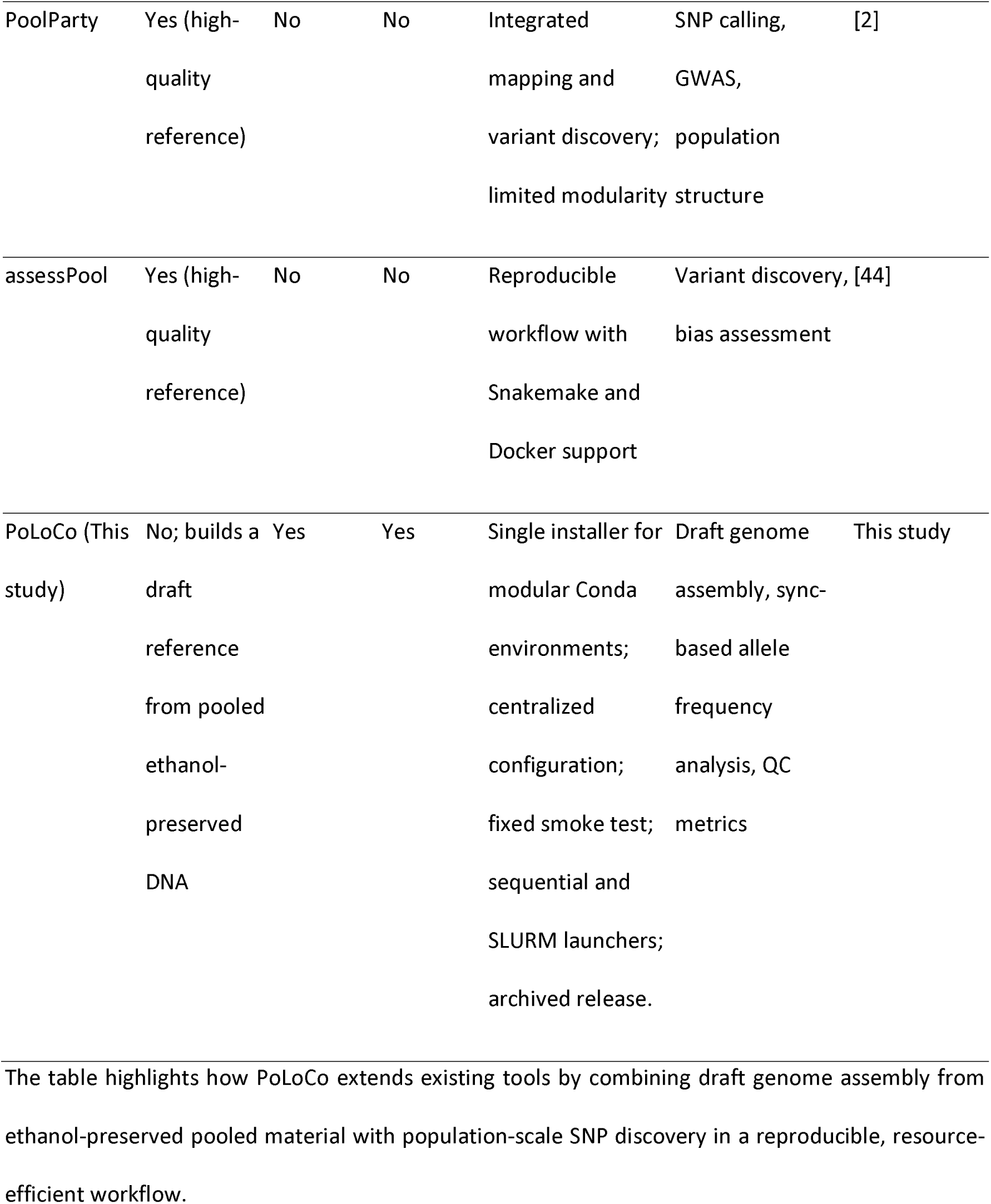
Comparison of common Pool-seq pipelines and PoLoCo.

### 3.3 Workflow performance and implications

The schematic representation of the general format of the workflow, including input samples, QC checkpoints, and parallel tracks, is illustrated in Figure 1. The PoLoCo workflow presented here combines raw read preprocessing, genome assembly, assembly validation, and population-scale allele-frequency analysis within a single reproducible workflow. Computational resource use and runtimes for the *E. nivalis* case-study execution were recorded on bwUniCluster 3.0 to provide practical information on the resources required for the major workflow stages. These records describe the computational scale of the present case study and are not intended as a controlled performance comparison with alternative Pool-seq software. No workflow step required more than 32 CPU cores or 128 GB of RAM, making the workflow very amenable to available infrastructure. Runtime and memory limits for each workflow step are provided in a brief format, summarized in Table 1. The full workflow, along with modular conda environments, SLURM scripts for each workflow step, and Python modules for quality control visualization, is publicly available in Zenodo [28]. The Zenodo-archived PoLoCo workflow package additionally includes centralized configuration, the Step 0 input check, a fixed end-to-end smoke test with defined expected outputs, sequential execution, and dependency-aware SLURM submission options. The optional final population-structure and spatial-connectivity module (Step 8) was also successfully executed using the complete 82-population case-study allele-frequency matrix, generating PCA outputs, pairwise genetic and geographic distances, spatial visualizations, and a 9,999-permutation Mantel test without manual reformatting of the PoLoCo allele-frequency matrix.

## 4. Discussion

The PoLoCo workflow presents a reproducible and scalable way of generating draft genomes and population-level genomic data from ethanol-preserved, non-model arthropods. It addresses challenges associated with degraded DNA in Pool-seq studies by integrating careful read filtering, assembly from degraded DNA, draft genome validation, sync-based allele-frequency analysis, and computational resources benchmarking. The workflow highlights key methodological considerations, practical limitations, and its potential broader applicability to other small-bodied specimens.

### 4.1. Generating genomic resources from ethanol-preserved material

The quality profiles for the draft genome assembly and pooled libraries highlight the feasibility of obtaining high-quality genomic data from ethanol-preserved Collembola specimens, as demonstrated by key standard QC metrics (Figure 2a-d). Quality metrics from all libraries fell within the expected ranges based on prior high-quality Illumina sequencing, indicating that even compromised material can produce reliable data. This indicates that ethanol-preserved specimens, often deemed unusable for genomics, can contribute to genome assemblies and population-level inference if developed with rigorous quality control and a reproducible protocol [14,45]. Comparable findings have recently been reported in studies showing that low-input or degraded DNA can still produce informative assemblies when processed with optimized pipelines [46]. In small arthropods, successful genomic sequencing has been reported from approximately 1–10 ng of degraded DNA in historical specimens and from as little as 5 ng in ethanol-preserved Collembola when appropriate low-input library-preparation strategies were used [39,46]. These examples indicate that workable DNA input is strongly dependent on preservation history, DNA integrity, and library-preparation chemistry rather than on a single universal concentration or fragment-length threshold.

### 4.2. Balancing fragmentation and completeness in draft assemblies

The draft genome assembly of *E. nivalis* showed low contiguity but high completeness, as indicated by fragmented assembly statistics in Table 2 and high BUSCO recovery in Table 3. The low contiguity, characterized by N50 of 3,617 bp, and the number of contigs is expected due to the degraded DNA and short reads used to sequence that assembly. Although short-read sequencing of degraded pooled material has clear limitations for assembly contiguity, the BUSCO analysis detected almost all conserved arthropod orthologs (97.7%), indicating that the vast majority of the functional gene set is present. The high duplicated-BUSCO fraction (46.3%), together with the larger assembly span (411.4 Mb versus 300.9 Mb for the published reference), is consistent with substantial assembly redundancy and may partly reflect uncollapsed alternative haplotypes arising from the pooled, heterozygous input. Thus, the high BUSCO completeness should be interpreted as broad gene-space recovery rather than as evidence of a nonredundant assembly. Haplotig purging was not applied to the present highly fragmented short-read assembly because distinguishing redundant haplotypes from genuine duplicated sequence would be uncertain under these conditions. Future applications with more contiguous assemblies and informative coverage profiles could evaluate haplotig-purging approaches to reduce redundancy. Whole-genome comparison with the published *E. nivalis* reference genome showed that the draft assembly recovered 97.9% of the reference genome fraction and had 62 misassemblies relative to the published assembly (Table 2). The complementary MUMmer4 dnadiff analysis was more restrictive: one-to-one aligned regions had an average identity of 87.96%, while aligned bases represented 38.65% of the published reference and 29.58% of the draft assembly. The QUAST genome-fraction statistic and the dnadiff aligned-base fractions therefore should not be interpreted as equivalent measures, because they are derived from different alignment summaries. The assembly comparison reflects several differences between the two genomic resources, including their source populations, preservation and sequencing histories, and assembly strategies. Because these factors were not experimentally separated in the present study, their individual contributions to the observed sequence and structural differences cannot be quantified. Besides, geographic population divergence has been noted in *Entomobrya*, including high degrees of cryptic diversity in the genus [47]. We therefore use the published genome primarily as an external reference for evaluating assembly correspondence and for testing how reference choice influences downstream Pool-seq results. Accordingly, PoLoCo is not intended to replace the more contiguous published *E. nivalis* reference genome (GCA_034695485.1). Instead, it provides a draft genome assembly derived from the ethanol-preserved specimens used in this study, and its high completeness, together with its observed mapping performance in the present case study, supported its use for SNP discovery and allele-frequency estimation. For Pool-seq-based population genomic analyses, assembly completeness is often more important than contiguity, provided that the reference retains sufficient genomic representation for reliable read mapping and SNP discovery [14]. Although high assembly completeness provides broad genomic representation, extensive fragmentation can still affect read placement, local SNP discovery, and allele-frequency estimation, particularly in repetitive or ambiguously mapped regions [16,17]. A project-specific reference may reduce sequence divergence from focal populations in some genomic regions, but this potential advantage must be balanced against mapping ambiguity caused by reference fragmentation and cannot be assumed to reduce reference bias universally. Competitive or multi-reference mapping approaches may provide an additional strategy for evaluating ambiguous alignments in highly fragmented or repeat-rich references, although such approaches were not implemented as a mandatory component of PoLoCo. The *E. nivalis* draft genome supplies population-specific genomic information suitable for SNP-based analyses within this project and can guide similar efforts in other understudied soil arthropods. The low contiguity of the draft assembly nevertheless limits analyses that depend on intact gene models or genomic context. Functional annotation, gene-level enrichment, and ecological interpretation of allele-frequency changes within fragmented genes should therefore preferentially use a more contiguous annotated reference where one is available.

### 4.3. Reference-dependent effects in pool-seq studies

Given the trade-off between assembly completeness and contiguity, reference genome choice can strongly influence SNP discovery and downstream population genomic inference [16,17]. The comparison of SNP discovery across the two reference genomes illustrates how reference choice can alter the composition of the resulting Pool-seq dataset. Although both the draft and published NCBI *E. nivalis* genomes enabled the discovery of genome-wide SNPs, the detailed measures indicated systematic differences that mattered for inference. The comparison of filtering outcomes across references showed that the draft genome yielded a larger number of loci after the minimum informative-population filter, but many of these loci were removed during minor allele frequency filtering and distance-based thinning, resulting in a substantially smaller final SNP set than that retained with the NCBI reference (Figure 4a). In contrast, the NCBI reference retained many more SNPs in the final dataset, indicating that reference choice can strongly influence the number of markers available for downstream analyses. At the same time, the draft-based dataset retained stronger population support per locus, with final SNPs more often represented across a greater number of the 82 pooled populations (Figure 4b). This pattern suggests that the project-specific draft genome provided a more conservative but more uniformly represented dataset for downstream analyses. Post-filtering minor allele frequency distributions also differed between references, with the draft-based dataset showing a stronger enrichment of low-frequency variants and the NCBI-based dataset showing a flatter retained minor allele frequency distribution (Figure 4c). Furthermore, the Ts/Tv ratios supported this difference in retained variant composition (Figure 4d). While the NCBI reference produced a higher Ts/Tv estimate than the draft genome, both values remained within the range reported for arthropod genomes [40–42]. Together, these results show that even when both references permit genome-wide SNP discovery, the final retained datasets can differ markedly in both size and composition. These results do not suggest that the fragmented draft genome is universally superior to the published reference. Rather, they show a clear trade-off between SNP yield and locus representation across populations. For Pool-seq studies aimed at population-level ecological inference, this trade-off is important because robust downstream analyses depend not only on marker number but also on consistent representation across populations. In this context, our draft genome generated by PoLoCo provides a project-specific reference that supported the focal study analysis, while the comparison with the published genome illustrates how strongly reference selection can shape final allele-frequency datasets.

### 4.4. Reproducibility, scalability, and broader applicability of PoLoCo

Our results indicate that PoLoCo is a viable, reproducible option for genomic studies on small, ethanol-preserved invertebrate taxa. In the *E. nivalis* case study, no workflow step was allocated more than 32 CPU cores or 128 GB RAM, indicating that the analysis can be implemented on standard high-performance computing infrastructure. Pool-seq sync generation and downstream allele-frequency matrix construction for the retained pooled libraries were performed efficiently on standard high-performance computing infrastructure, and the assembly of a draft genome together with BUSCO validation took roughly 48 hours. The recorded runtimes and resource allocations for the major workflow stages are summarized in Table 1 and provide practical guidance for planning comparable analyses. These benchmarks describe the computational scale of the present case study rather than a controlled performance comparison with alternative Pool-seq software. A major strength of PoLoCo is its focus on reproducibility. The workflow is provided as an open-source Zenodo archive with modular conda environments, SLURM scripts, and automated QC visualization. The modular structure of the PoLoCo Zenodo archive allows users to either execute the workflow end-to-end or run individual components independently, enabling flexible adaptation to different datasets, organisms, and research objectives. The fixed smoke-test dataset and case-study input utilities further allow users to verify installation and core workflow behavior without first preparing a complete independent dataset. This approach is consistent with established recommendations for practical computational reproducibility in the life sciences [48]. Together, these features make the computational steps, software environments, and configurable parameters explicit from raw-read trimming and sync-based SNP filtering through assembly validation. Furthermore, the add-on design makes the package usable for researchers of all computational skills and applicable to other taxa. Thus, it is well-aligned with current recommendations to standardize open pipelines for pool-seq analyses [17,44]. However, in contrast to these existing pipelines, PoLoCo includes genome assembly from ethanol-preserved pools for downstream sync-based allele-frequency analysis (Table 4), allowing project-specific reference generation and extending Pool-seq analyses to study systems for which obtaining high-quality DNA is difficult.

To place these features in context, we next compared PoLoCo with existing Pool-seq analysis frameworks. Over the past decade, several Pool-seq pipelines have been developed to enable efficient and scalable pooled allele-frequency analysis, including PoPoolation2 [4], SNAPE-pooled [18], ANGSD [43], PoolParty [2], and assessPool [44]. These frameworks provide different approaches to pooled variant or allele-frequency analysis, including count-based and genotype-likelihood-based inference, yet they all rely on an available reference genome to perform downstream analyses. In contrast, PoLoCo provides a unified draft genome assembly, reference validation, and a population-level sync-based allele-frequency analysis process within a single reproducible workflow. This integration is particularly valuable when working with degraded or field-preserved material, enabling creation of project-specific references while allowing the consequences of reference choice to be evaluated explicitly in the resulting allele-frequency datasets.

### 4.5. Implications for biodiversity and ecological genomics

Utilizing a reliable short-read assembly strategy combined with Pool-seq allele-frequency analysis, PoLoCo applies the methodological toolbox to non-model invertebrates. Beyond this proof-of-concept in *E. nivalis*, the workflow is broadly applicable to other small-bodied taxa that remain underrepresented in current genomic resources despite growing biodiversity genome initiatives, often because suitable material for reference generation is difficult to obtain from field-collected specimens [8,10]. Natural-history collections also contain large numbers of preserved specimens that are increasingly being considered for genomic research, although DNA quantity and integrity can vary substantially with specimen age and preservation method [45,50]. Advances in recovering genomic data from such material are expanding opportunities for ecological and evolutionary genomics beyond freshly collected specimens. The present PoLoCo case study specifically evaluates ethanol-preserved *E. nivalis* material; applications to substantially older material or other preservatives would require input-quality assessment and project-specific adjustment of the configurable workflow parameters. Likewise, the 6–10-individual population pools used here represent the case-study design rather than a fixed PoLoCo requirement; studies using different pool sizes can retain the same workflow structure while adjusting coverage and population-representation settings and using pool-size-aware estimators where required for downstream population-genetic analyses. This framework complements larger biodiversity genomics initiatives focused on reproducibility and accessibility of genomic resources for non-model taxa. For instance, large-scale community-based initiatives like i5K Arthropod Genomes [7] and MetaInvert [8] show the importance of community-based resources needed for invertebrates, but they primarily rely on freshly collected specimens and high-quality DNA. In contrast, PoLoCo provides an organizational framework for studies in which specimens are ethanol-preserved and collected under field-based circumstances, thereby extending genomic analyses to situations where ideal material is not available. Together, our results show that tailored workflows can help address practical obstacles in biodiversity genomics, particularly for non-model arthropods preserved under field conditions.

## 5. Conclusion

The present study provides a reproducible pooled low-coverage workflow for draft genome assembly and allele-frequency analysis from ethanol-preserved invertebrates, integrating draft genome construction, reference validation, and population-level Pool-seq analysis. Taking *E. nivalis* as a case study, we demonstrate that pooled short-read sequencing of highly degraded samples can produce a near-complete genome and can provide large-scale estimates of allele frequencies across 82 retained pooled populations. Benchmarking shows that PoLoCo operates efficiently on standard high-performance computing resources, with all code and environments openly available. PoLoCo is designed as a programmable and adaptable workflow for ethanol-preserved, non-model organisms, with project-specific adjustment of input-quality assessment and filtering parameters where preservation history, DNA quality, or pool sizes differ from the present case study. It thus provides a practical and transferable framework for generating and validating draft genome assemblies, and for supporting downstream population genomic inference, in ecological and evolutionary studies where high-quality DNA or long-read sequencing is not feasible.

## Author Contributions

MJS: Conceptualization, Methodology, Formal analysis, Investigation, Data curation, Visualization, Writing - original draft. GS: Conceptualization, Funding acquisition, Supervision, Writing - review & editing. JCG: Investigation, Validation, Supervision, Writing - review & editing. All authors read and approved the final manuscript.

## Acknowledgements

We are grateful to Forst Baden-Württemberg (ForstBW) for enabling and facilitating the fieldwork without their excellent support; our study would not have been possible. Our thanks go to the ConFoBi team for their dedicated involvement and valuable contributions to this project, especially Dr. Johannes Penner for his assistance with the fieldwork, sorting the specimens, and storing them in our facilities. We also thank Dr. Michael Wohlwend and Susanne Hartmann for coordinating the administrative matters. We also thank the anonymous reviewers for their careful and constructive comments, which helped us substantially improve the manuscript and the PoLoCo workflow. We gratefully acknowledge the support of the German Science Foundation (DFG), Research Training Group ConFoBi (GRK 2123/1 TPX). We also acknowledge support by the state of Baden-Württemberg through bwHPC for this project.

## Data Availability

The raw sequencing reads supporting this article are available in the European Nucleotide Archive (ENA) under BioProject accession PRJEB111482 [37]. The draft genome assembly, BUSCO outputs, QUAST reports, and supplementary datasets are available in Figshare [38]. The PoLoCo workflow, source code, conda environments, SLURM scripts, and QC utilities are available in Zenodo [28].

## Availability of Source Code and Requirements

Project name: PoLoCo

Project home page: GitHub repository for PoLoCo-workflow [49]

Archived version: PoLoCo-workflow v1.1.0, Zenodo [28]

Operating systems: Linux; tested on bwUniCluster 3.0 HPC systems

Programming language: Bash, Python, R

Other requirements: Conda; SLURM optional for HPC batch execution

License: MIT License

RRID: not applicable

bio.tools ID: not applicable

